# Niche Partitioning Between Coastal and Offshore Shelf Waters Results in Differential Expression of Alkane and PAH Catabolic Pathways

**DOI:** 10.1101/2020.03.16.994715

**Authors:** Shawn M. Doyle, Genmei Lin, Maya Morales-McDevitt, Terry L. Wade, Antonietta Quigg, Jason B. Sylvan

## Abstract

Marine oil spills can impact both coastal and offshore marine environments, but little information is available on how the microbial response to oil and dispersants might differ between these biomes. Here we describe the compositional and functional response of microbial communities to different concentrations of oil and chemically dispersed oil in coastal and offshore surface waters from the Texas-Louisiana continental shelf. Using a combination of analytical chemistry, 16S rRNA amplicon, and metatranscriptomic sequencing, we provide a broad, comparative overview of the ecological response of hydrocarbon degrading bacteria and their expression of hydrocarbon degrading genes in marine surface waters over time between two oceanic biomes. We found evidence for the existence of different ecotypes of several commonly described hydrocarbon degrading bacterial taxa which behaved differentially in coastal and offshore shelf waters despite being exposed to similar concentrations of oil, dispersants, and nutrients. This resulted in the differential expression of catabolic pathways for n-alkanes and polycyclic aromatic hydrocarbons (PAH)—the two major categories of compounds found in crude oil—with preferential expression of n-alkane degradation genes in coastal waters while offshore microbial communities trended more towards the expression of PAH degradation genes. This was unexpected as it contrasts with the generally held view that n-alkanes, being more labile, are attacked before the more refractory PAHs. Collectively, our results provide new insights into the existence and potential consequences of niche partitioning of hydrocarbon degrading taxa between neighboring marine environments.

**IMPORTANCE:** In the wake of the Deepwater Horizon oil spill, the taxonomic response of marine microbial communities to oil and dispersants has been extensively studied. However, relatively few studies on the functional response of these microbial communities have been reported, especially in a longitudinal fashion. Moreover, despite the fact that marine oil spills typically impact thousands of square kilometers of both coastal and offshore marine environments, little information is available on how the microbial response to oil and dispersants might differ between these biomes. The results of this study help fill this critical knowledge gap and provide valuable insight into how oil spill response efforts, such as chemically dispersing oil, may have differing effects in neighboring coastal and offshore marine environments.

## INTRODUCTION

Crude oils are complex mixtures of thousands of different compounds, ranging from saturated alkanes and aromatic hydrocarbons to complex, heteroatom-containing resins and asphaltenes (1). While some of these compounds are recalcitrant to degradation, many can be utilized as a carbon or energy source by microorganisms (2). Microbial biodegradation is one of the primary means by which both natural and anthropogenic oil releases are bioremediated in nature. However, many crude oil hydrocarbons are poorly soluble, which lowers their availability to microorganisms and limits biodegradation rates. As such, chemical dispersants have been used after marine oil spills to disperse oil into the water column with the aim of dramatically increasing the surface area for microbial attack (3). Despite this, studies have reported conflicting results on whether chemical dispersants improve or suppress oil biodegradation rates by microbes (4–8).

The Deepwater Horizon (DwH) oil spill involved the release of ∼780,000 m^3^ crude oil into the Gulf of Mexico after which 7000 m^3^ chemical dispersants were used during the spill response effort (9, 10). The magnitude of the spill led to a surge of research into the microbial ecology of marine oil spills and dispersant usage both in deep-sea habitats (11–15) and shoreline environments such as beaches (16–19) and salt marshes (20, 21). However, much less information is available on the responses of microbial communities in surface waters on the continental shelf (22). These areas in the Gulf of Mexico range from river-dominated, nutrient rich waters along the coasts to near oligotrophic waters on the continental shelf edge. Oil spills can affect many thousands of square kilometers of both open ocean and coastal habitats, and the microbial communities which inhabit these respective marine biomes can differ substantially in composition, structure, and metabolic potential (23, 24). As a result, natural biodegradation of oil spills likely varies considerably across different shelf water regimes. Understanding how microbial communities inhabiting these different ocean biomes respond to oil and dispersants consequently represents an important research need.

To better define the ecological and functional response of marine microorganisms to oil, we conducted mesocosm experiments using coastal seawater collected from the continental shelf (herein referred to as Coastal) and offshore waters on the edge of the shelf (herein referred to as Offshore) of the northwestern Gulf of Mexico (Figure S1, Table S1) and amended them with oil or chemically dispersed oil. The responses of the microbial communities were subsequently followed over time using cell counts, 16S rRNA amplicon and metatranscriptomic sequencing. This analysis allowed us to identify which groups of bacteria responded to oil with and without dispersant and also characterize the functional response of hydrocarbon-degrading microbes within the communities. We found evidence for the existence of different ecotypes—amplicon sequence variants (ASVs) belonging to the same genus occupying the same niche—of several well-known hydrocarbon degrading bacterial taxa which behaved differentially in coastal and offshore shelf waters. Our results revealed that these ecotype differences between the coastal and offshore communities resulted in differential expression of alkane and PAH catabolic pathways between these oceanic biomes and that the response to dispersants was variable by location.

## RESULTS

Four treatments were prepared in triplicate for each experiment: [1] “Control”, containing only seawater, [2] Water Accommodated Fraction or “WAF”, containing seawater amended with the fraction of oil accommodated after physical mixing alone, [3] Chemically-Enhanced WAF or “CEWAF”, seawater amended with the fraction of oil accommodated after dispersal with Corexit, and [4] Diluted CEWAF or “DCEWAF”, a 1:10 dilution of the CEWAF treatment. Because the concentration of oil in the CEWAF treatment was so high, we included the DCEWAF treatment in order to observe the effects of a chemically dispersed oil, but at a concentration more commonly encountered at sea after an oil spill (25).

### Hydrocarbon Chemistry

We tracked total alkane and PAH concentrations over time using a combination of GC/FID and GC/MS, respectively (Figure 1, Table S2). In the Control treatments of both experiments, these concentrations were <5 μg L^-1^ for both classes of compounds with one exception: the seawater used in the Coastal experiment contained a variable concentration of n-alkanes, ranging from 3.2 to 24 μg L^-1^, with an average odd/even ratio of 1.3 indicating a mixture of biogenic and petroleum sources. The biogenic n-alkanes appear to be both marine (e.g. n-C17) and terrestrial odd chained (e.g. n-C27) n-alkanes. In the WAF treatments of both experiments, alkane concentrations were also very low and did not differ substantially from those observed in the Controls. Initial PAH concentrations were elevated to ∼50 μg L^-1^ but rapidly dropped after 24 hours, likely due to the evaporation of volatile PAHs entrained into oil droplets during WAF production. In the DCEWAF and CEWAF treatments, chemically dispersing the oil increased initial alkane concentrations to ∼325 μg L^-1^ and ∼3000 μg L^-1^, respectively. Initial PAH concentrations within these two dispersed oil treatments were also increased, but more modestly, to ∼100 μg L^-1^ and ∼400 μg L^-1^, respectively.

**Figure 1.**
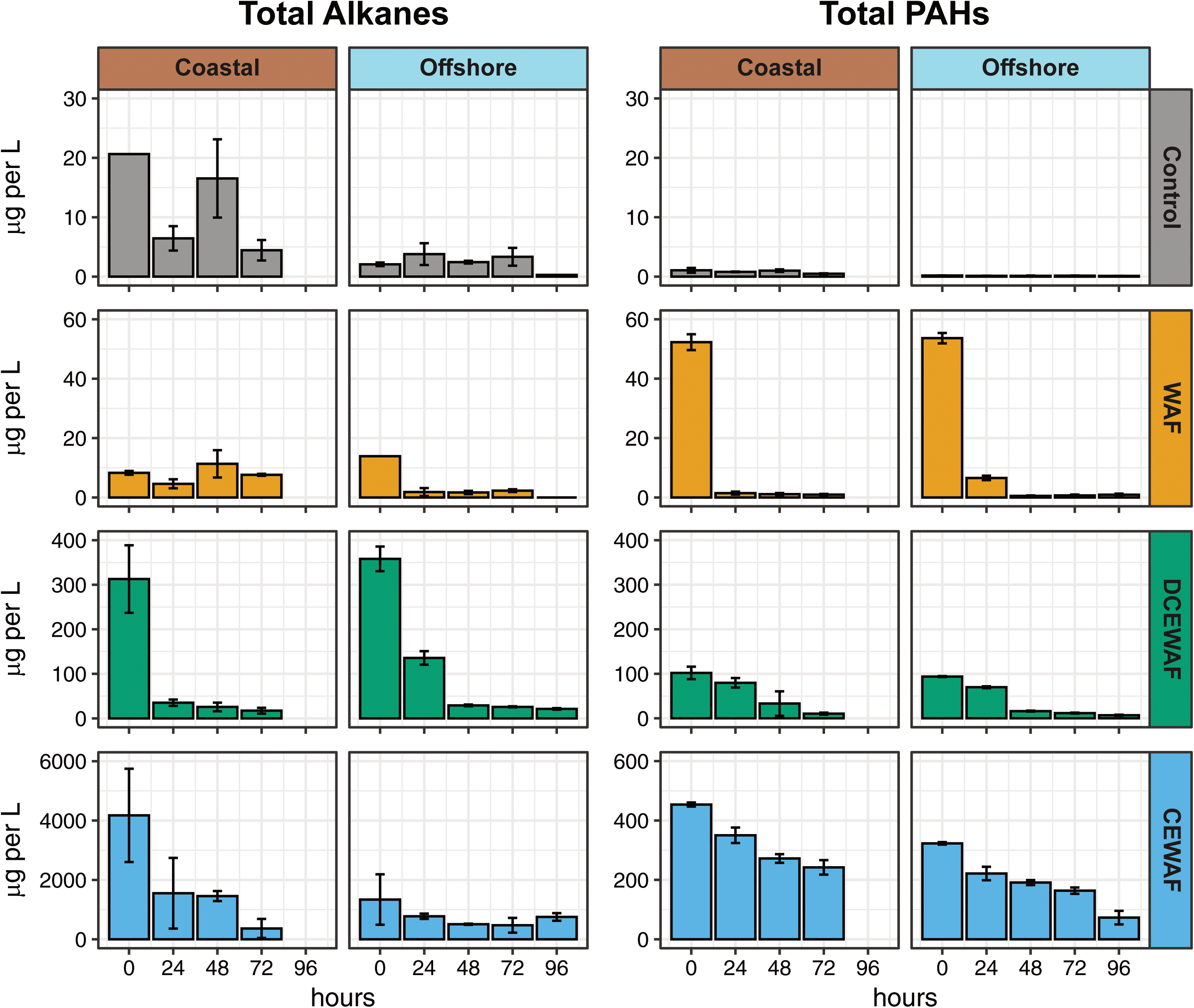
Measured concentrations (µg/L) of total alkanes and total PAHs over time in both mesocosm experiments. Error bars denote standard deviation.

### Microbial cell abundance

Average cell abundances (Figure S2) were typical for seawater, ∼10^6^ cells mL^-1^ (26, 27). The average abundance in the Coastal experiment (2.4±0.7 ×10^6^ cells mL^-1^) was twofold that in the Offshore experiment (1.3±0.5 ×10^6^ cells mL^-1^). Between treatments, cell abundances were ∼1.7-fold higher in the Coastal CEWAF treatment [*F*(3,20)=11.5, *p*<0.001] than in the Control, WAF, and DCEWAF treatments, where cell abundances were similar [*F*(2,18)=1.22, *p*=0.32]. In the Offshore experiment, cell abundances were instead slightly lower (1.6-fold) in the Control treatment [*F*(3,32)=3.03, *p*=0.04], while those in the WAF, DCEWAF, and CEWAF remained similar [*F*(2,24)=0.78, *p*=0.47]. Using simple linear regression models, we found cell abundances increased over time in the Offshore DCEWAF, Offshore CEWAF, and the Coastal CEWAF while those in the Control and WAF treatments of both experiments generally decreased (Table S3). However, in all cases, these changes were relatively small (±0.6% per hour).

### Microbial community composition

We profiled the composition and structure of the microbial communities using 16S rRNA amplicon sequencing (Figure 2). NMDS ordination of Bray-Curtis dissimilarities (BC-D) confirmed the Coastal and Offshore experiments harbored distinct microbiomes (Figure S3). In the Offshore experiment, samples formed four distinct clusters organized sequentially by the four treatments (Figure S4A and S4E). Relative to the Control treatments, communities exposed to WAF were the most similar (BC-D: 0.30 ± 0.09; avg ± SD), followed by DCEWAF (0.47 ± 0.13), and finally CEWAF (0.59 ± 0.14). A similar pattern was observed in the Coastal experiment. Communities exposed to CEWAF were the most distinct from the Controls (0.82 ± 0.10), followed by the DCEWAF (0.58 ± 0.14) and WAF (0.39 ± 0.09), respectively. Differences in community structure observed at the first time point (0 hours) indicate some shifts in microbiome composition and structure occurred during the preparation of WAF, DCEWAF, and CEWAF (∼24 hours).

**Figure 2.**
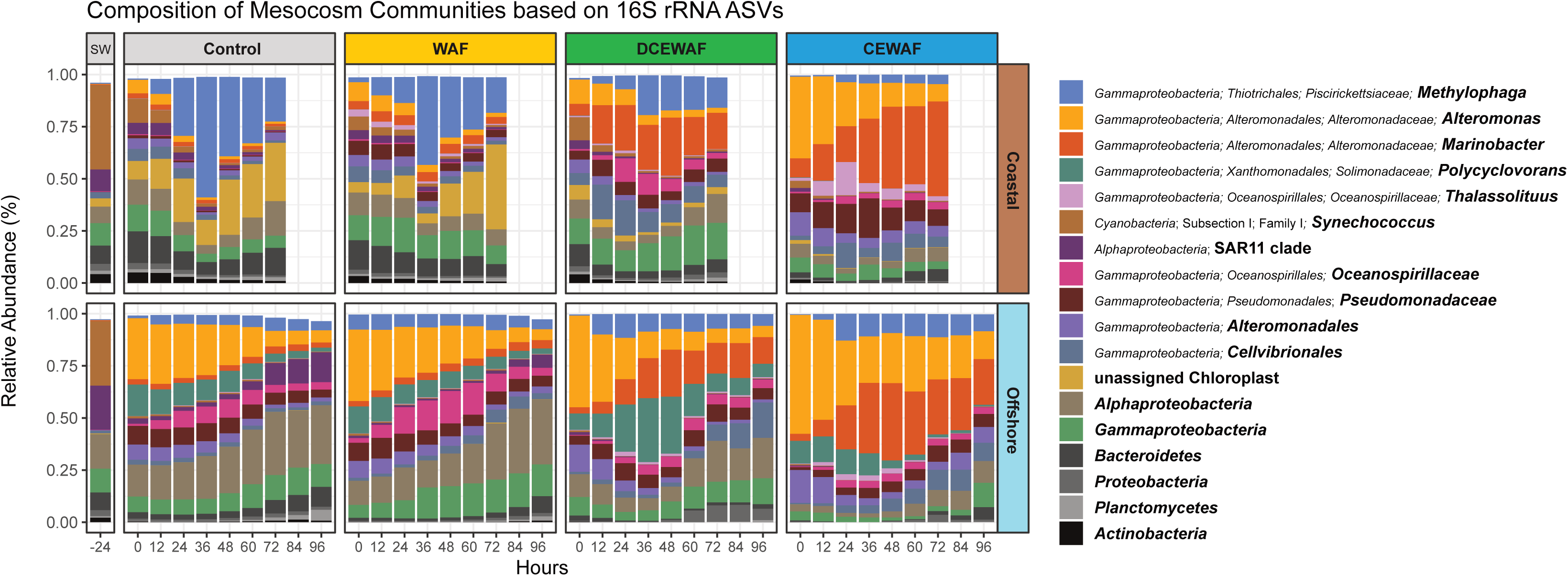
Relative abundances of abundant microbial lineages observed in each experiment. Each bar is the average of triplicate treatments. Community composition in the original seawater samples collected for each experiment are shown under category “SW”. The plot was built to display the highest resolution classification for the most abundant taxa. The plot was constructed as follows: First, ASVs were clustered at the genus level and any genera having a relative abundance ≥15% in at least one of the samples were plotted. This procedure was subsequently repeated with the remaining unplotted ASVs at the taxonomic levels of family, order, class and phylum. Any remaining rare ASVs left after this procedure were not plotted.

In total, we detected 2880 ASVs across both experiments. Compositionally, *Gammaproteobacteria* was the most abundant group observed, constituting an average of 77.4% and 67.0% of the overall communities within the Offshore and Coastal experiments, respectively (Figure 2, Figure S5). Within this class, dominant taxa included *Alteromonadales* (*Alteromonas*, *Marinobacter*, and *Aestuariibacter*), *Thiotrichales* (*Methylophaga* and *Cycloclasticus*), *Oceanospirillales* (*Alcanivorax*, *Oleibacter*, and *Thalassolituus*), and *Xanthomonadales* (*Polycyclovorans*) (Figure S6).

### Identification of oil-enriched ecotypes

ASVs from the same genus often varied in how they responded to oil or dispersed oil (Table S4). These differences occurred both between treatments and between the two experiments. In order to more accurately define potential ecotype responses, we used redundancy analysis (RDA) modeling to narrow our dataset and identify only those ASVs which were directly enriched by oil within the WAF, DCEWAF, and CEWAF treatments. In each model, variables for both time and initial oil concentration contributed significantly (*p*<0.001) to the RDA models (Figure S4, B-D and F-H). Using scalar projections of ASVs onto the *Oil* vector in each model, we identified 30 oil-enriched ASVs between the two experiments, 8 of which were identified in both the offshore and coastal shelf water mesocosms (Figure 3).

**Figure 3.**
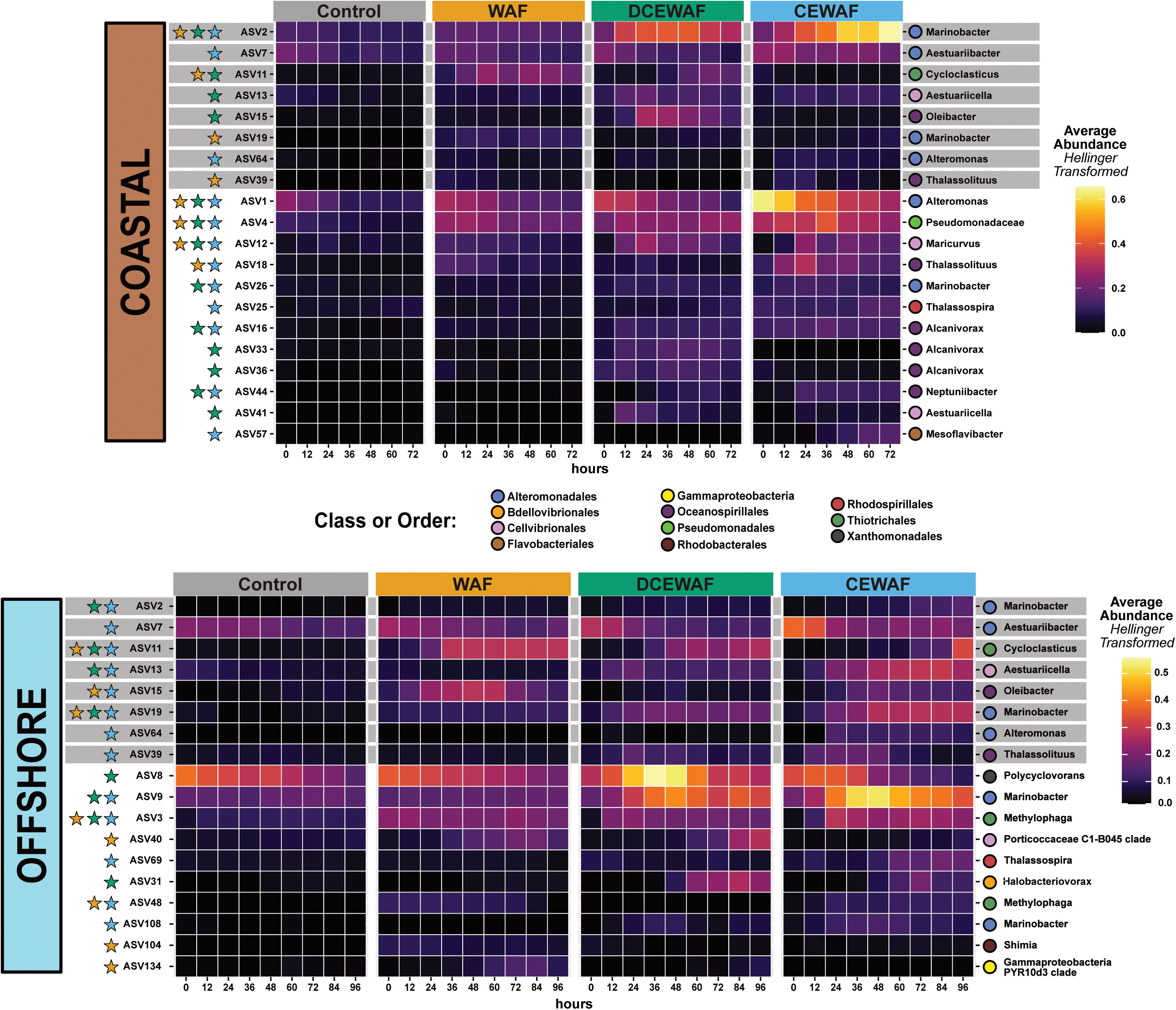
Heatmap displaying the relative abundances of the 30 ASVs identified as being oil-enriched using the RDA models. ASVs identified in both experiments are highlighted with a gray bar. Colored stars on the left side indicate the treatment(s) in which an ASV’s relative abundance over time was significantly enriched versus the Control. ASV read counts were averaged among replicates and transformed with a Hellinger transformation in order to facilitate a clearer visual comparison between abundant and sparse taxa.

The response of many oil-enriched ASVs to each treatment was notably different between the two experiments. For example, we found three *Alcanivorax*-related ASVs were only enriched by oil in the Coastal experiment; in the Offshore experiment these were either very rare (e.g. ASV33 and ASV36) or were not significantly more abundant in the oil amended treatments versus the Control (ASV16). Likewise, ASV8 (*Polycyclovorans*)—an obligate hydrocarbon degrading taxa which inhabits the phycosphere of marine diatoms and dinoflagellates (28)—was present in both experiments, but was enriched only in the Offshore DCEWAF treatment. We also found several ecotypes which exhibited differential responses to oil between the two experiments. For example, ASV2 (*Marinobacter*) was highly abundant in the Coastal experiment (11.9%; average relative abundance) where it was enriched in all three oil-amended treatments. In contrast, ASV2 was comparatively sparse Offshore (0.3%) and was instead only enriched in the DCEWAF and CEWAF. Meanwhile, another *Marinobacter*-related ASV (ASV19), was abundant in the Offshore WAF, DCEWAF, and CEWAF treatments, but was restricted to only the WAF treatment in the Coastal experiment. Lastly, ASV11 (*Cycloclasticus*) increased in relative abundance over time similarly in both the Coastal and Offshore WAF and DCEWAF treatments, but had an opposite response to CEWAF between the two experiments. In the Offshore CEWAF, ASV11 exhibited a large bloom from ≤0.1% to 10.3% relative abundance over the course of the experiment while in the Coastal CEWAF, it instead decreased over time from 0.5% to 0.1%.

### Identification of hydrocarbon degradation gene transcripts

We parsed the metatranscriptome datasets to identify and quantify the expression of hydrocarbon degradation gene transcripts within the mesocosms. A GHOSTX query of the KEGG database identified 6195 transcripts affiliated with hydrocarbon degradation genes. An additional 163 transcripts were identified as alkane hydroxylating P450 genes from the CYP153 family through a custom BLASTP search. We compared the abundance of these hydrocarbon degradation gene transcripts between each oil-amended treatment and its respective control treatment and used an outlier analysis to estimate the number of upregulated genes at each time point (Figures S7 and S8). Within both experiments, this number was generally higher in the DCEWAF and CEWAF treatments versus the WAF treatments. We also observed that the number of upregulated genes increased overtime in dispersant-amended treatments, but decreased overtime in WAF.

We next sought to test if the amount of upregulation among these upregulated hydrocarbon degradation genes was affected by treatment. Highly expressed hydrocarbon degradation gene transcripts were on average 10.4-fold more abundant in the oil-amended treatments than in the Controls. However, using a factorial ANOVA test with pairwise comparisons of the oil-amended treatments, we found only a single significant difference between the Coastal WAF and CEWAF, which was small (1.2-fold; *t*(4)=-3.44, p=0.03). In other words, the expression of upregulated hydrocarbon degradation genes was higher in the WAF, DCEWAF, and CEWAF treatments over the Controls, but by about the same amount per oil-amended treatment (Figure S9). This suggests a high degree of functional redundancy among community members and indicates that the expression of hydrocarbon degradation genes was primarily structured by microbial community turnover—as we observed in our 16S rRNA amplicon dataset.

### Coastal versus offshore expression of n-alkane and PAH degradation pathways

To compare differences in the expression of hydrocarbon catabolic pathways in our experiments, we consolidated the identified gene transcripts into four metabolic categories: [1] n-alkane activation, [2] beta-oxidation of fatty acids, [3] ring-hydroxylating/cleaving dioxygenases, and [4] PAH degradation (Figure 4). Alkane degradation begins with terminal oxidation of the substrate to a primary alcohol with a hydroxylase or monoxygenase enzyme (e.g. AlkB). The resulting alcohol is further oxidized to a fatty acid by alcohol and aldehyde dehydrogenases before entering the beta oxidation pathway (29). Likewise, PAH degradation typically begins with ring hydroxylation and subsequent cleavage with dioxygenase enzymes (30), the products of which are then channeled through various complex, multi-step catabolic pathways (31). Hence, these four categories together represent the activation [1+3] and subsequent degradation [2+4] pathways for saturated hydrocarbons (i.e. linear and cyclo-alkanes) and PAHs, respectively, to central metabolism intermediates.

**Figure 4.**
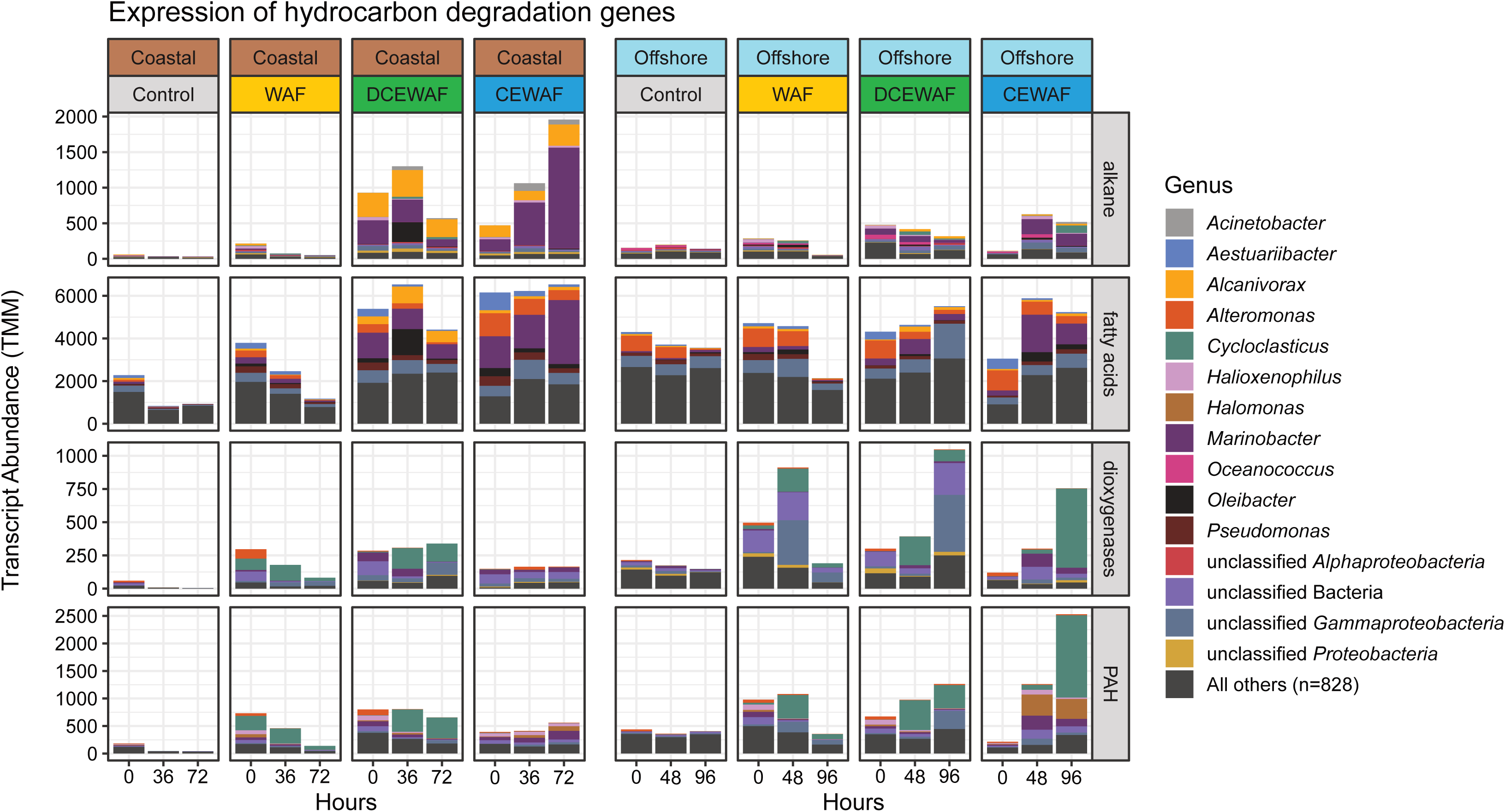
Coastal and offshore microbial communities exhibited differential expression of alkane and PAH catabolic pathways after exposure to oil. Taxonomic affiliations and abundances of hydrocarbon degradation gene transcripts are shown within four metabolic categories. Y-axes are individually scaled for each category (row). Transcripts belonging to minor taxa (<2% total abundance) were grouped together into the “All others” category.

Alkane activation gene transcripts were observed primarily in the DCEWAF and CEWAF treatments of the Coastal experiment where they were produced almost entirely by *Marinobacter* and *Alcanivorax* (Figure 4). In comparison, expression of alkane degradation genes in the Coastal WAF and Control treatments was very low (∼28 TPM; transcripts per million transcripts). This was consistent with the very low total alkane concentrations we measured in these two treatments (< 14 ug/L; Figure 1). In the Offshore experiment, expression of genes involved with alkane activation was instead comparatively low across all four treatments.

Expression of fatty acid beta-oxidation genes was the highest of the four metabolic categories. We observed transcripts from a wide range of taxa, reflecting the nearly universal taxonomic distribution of this pathway (32). In the Coastal experiment, highest expression levels were again observed in the DCEWAF and CEWAF with comparatively lower expression levels in the WAF and Control treatments. Similar to the alkane activation genes, fatty acid beta-oxidation genes in the Coastal experiment were expressed in large part by *Marinobacter* and *Alcanivorax* (Figure 4). However, significant expression of *Alteromonas*, *Aestuariibacter*, *Oleibacter*, and *Pseudomonas-*assigned transcripts were observed in this category as well. In the Offshore experiment, abundant transcripts were detected from *Alteromonas* in all four treatments, while those from *Marinobacter, Aestuaribacter* and *Alcanivorax* were mainly observed in DCEWAF and CEWAF.

In contrast to the alkane degradation categories, overall expression for ring cleavage/hydroxylation dioxygenase and PAH degradation genes was significantly higher [*t*(2760)=6.28, p<0.001] in the Offshore experiment (Figure 5). *Cycloclasticus*, a genus of obligate PAH degraders (33), were the primary source of detected ring cleavage/hydroxylation dioxygenase gene transcripts (∼27% of all transcripts in these two categories) across both experiments. These *Cycloclasticus*-derived dioxygenase gene transcripts were observed in every sample except the Controls and the Coastal CEWAF. This pattern was also observed in our 16S rRNA amplicon dataset wherein the relative abundance patterns of ASV11 (*Cycloclasticus*) was substantially lower in the Coastal CEWAF treatments (Figure 3). Other taxa to which a significant portion of dioxygenase and PAH degradation gene transcripts were assigned to were *Halomonas*, *Halioxenophilus*, unclassified *Gammaproteobacteria*, and unclassified Bacteria (Figure 4).

**Figure 5.**
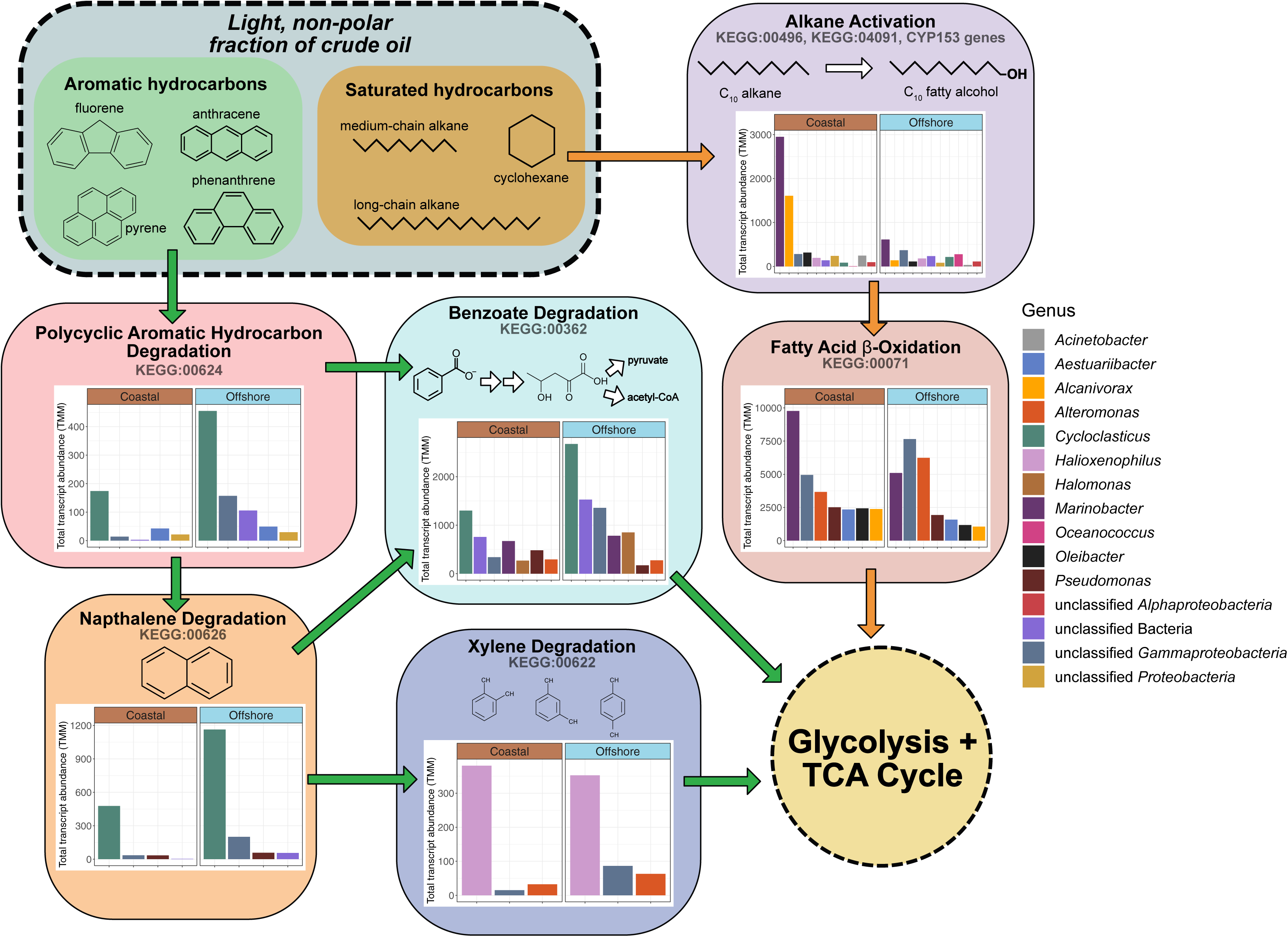
Summary overview of the taxonomic distribution and summed abundances, between all four treatments, of expressed hydrocarbon degradation genes involved in the transformation of n-alkanes (orange arrows) and PAHs (green arrows) to central metabolism intermediates.

In both experiments, the alpha diversity of taxa expressing alkane degradation genes decreased as oil concentrations increased (Control > WAF > DCEWAF > CEWAF) (Figure 6). In the categories for PAH degradation, we also observed a drop in alpha diversity when oil was present, but the difference between the three oil-amended treatments was minimal. These patterns indicate that as oil concentrations increase, a smaller subset of hydrocarbon degrading taxa are selected for and became functionally dominant.

**Figure 6.**
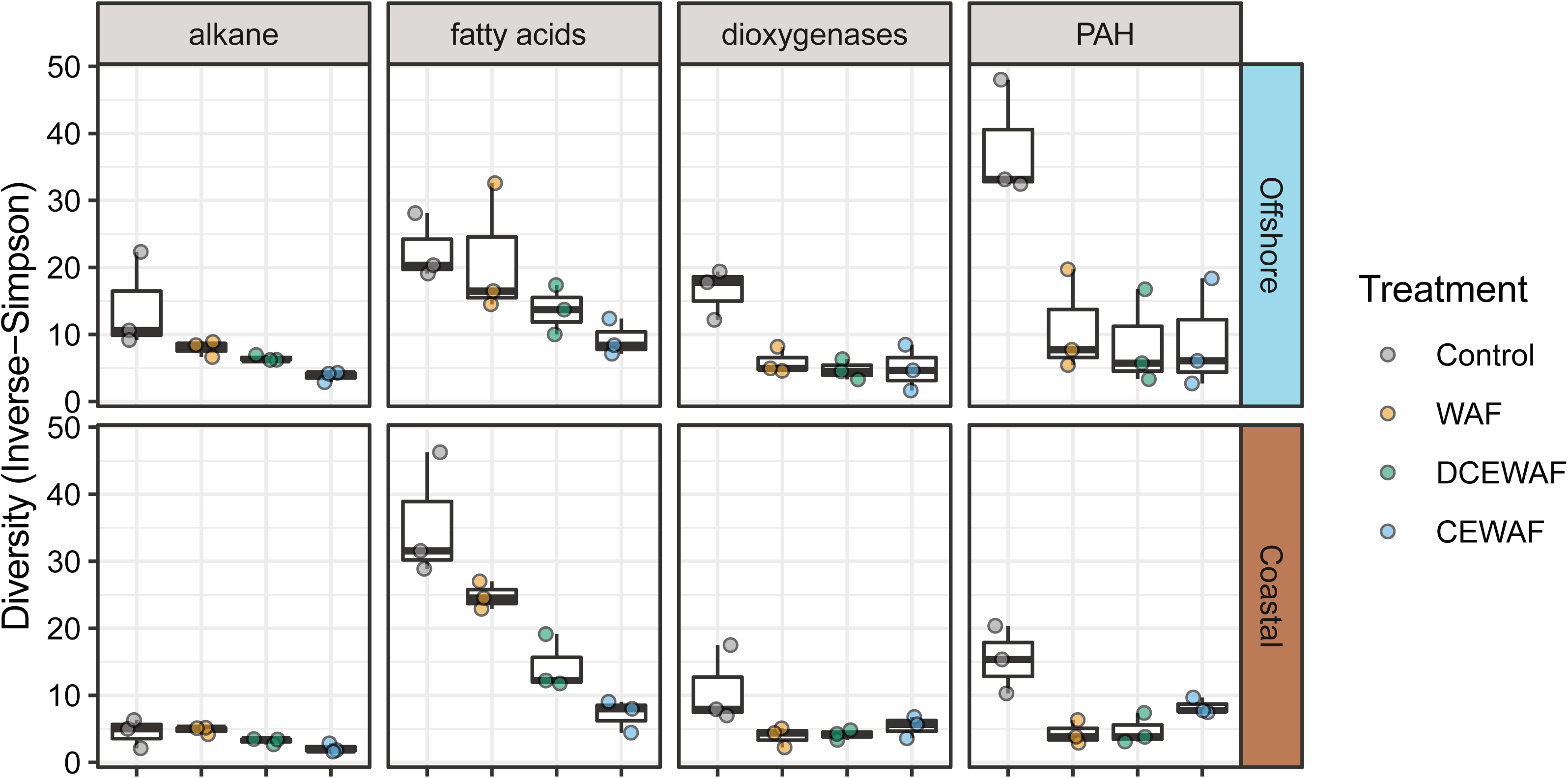
Alpha diversity of hydrocarbon degradation gene transcripts within the four metabolic categories. Boxplots display variation between time points for each treatment. Transcripts within each category were taxonomically classified using KAIJU (minimum score 65) and then clustered by genus before calculating Inverse-Simpson indices, a metric for effective species number (i.e. the number of equally common taxa).

## DISCUSSION

In this study, we found initial oil concentration strongly structured the microbial communities. Many of the identified bacteria belong to hydrocarbon degrading taxa which are known to bloom in seawater during oil spills or near natural oil seeps (2, 34, 35). However, we also found evidence that many of these taxa were composed of ecotypes which responded remarkably different to oil and dispersants in coastal and offshore shelf waters. These differences were reflected in the expression of hydrocarbon degrading genes, where we observed unexpected differences in alkane and PAH degradation pathways between the two oceanographic zones. Interestingly, patterns in the taxonomic diversity of expressed hydrocarbon degradation genes did not vary substantially between environments, indicating that some aspects of the response of microbial communities in marine surface waters to oil or chemically dispersed oil exposure may be common between locales. Overall, these findings highlight that natural remediation processes such as microbial biodegradation are not uniform across different ocean biomes (36) and reinforce the need for a better understanding of the spatial and temporal variances in natural oil remediation processes.

### Ecotype Dynamics

Following the distinct clustering we observed in the NMDS ordinations, many of the oil-enriched taxa (e.g. *Marinobacter*, *Alteromonas*, *Alcanivorax*, and *Polycyclovorans*) appeared to bloom in response to increasing oil concentrations, while others such as *Methylophaga* appeared to be either inhibited or outcompeted by other organisms with larger relative abundances. *Alteromonas* and *Aestuaribacter* were identified as oil-enriched in our 16S rRNA dataset (Figure 3) and have been repeatedly identified in laboratory and field based 16S rRNA surveys of marine oil spills (2, 37–40). In our metatranscriptomes, these two taxa were almost exclusively responsible for expression of fatty acid oxidation genes. This suggests that these two taxa were primarily secondary alkane degraders relying on other community members to activate alkanes (41). *Halomonas* were detected in the CEWAF treatments, but were not abundant elsewhere. It was not one of the ASVs enriched by oil, and is not known for PAH degradation, so it may also have responded to secondary metabolites in the CEWAF treatments.

In our experiments, we found three *Alcanivorax*-related ASVs that were enriched by oil. Two of these, ASV33 and ASV36, differed by only a single nucleotide position (Table S5) and had similar responses to the three coastal shelf water oil treatments: they grew well in the DCEWAF but remained relatively rare in the WAF and CEWAF. As such, these two ASVs probably represent a single ecotype of *Alcanivorax* which grows optimally at moderate oil concentrations, but only in coastal shelf waters, as they remained rare (≤0.7%) in all Offshore treatments. The third ASV (ASV16) differed from the other two ASVs at nine nucleotide positions and grew in not only the Coastal DCEWAF but the Coastal CEWAF treatment as well. Because *Alcanivorax* genomes typically contain 2 or 3 copies of the 16S rRNA gene (42), we sought to test if ASV33 and ASV36 might represent polymorphic 16S rRNA gene copies from a single *Alcanivorax* species. If this was the case, the relative abundance ratio between these two ASVs would be consistent across all samples. However, a one-way ANOVA found this was not the case [*F*(65,129)=2.45, p<0.001], indicating ASV33 and ASV36 are likely from separate from very closely related species of *Alcanivorax*. *Halioxenophilus* are known to degrade xylene (43), but this newly discovered genus has not been reported to degrade PAHs in an environmental context previously. In our study, PAH degradation gene transcripts assigned to *Halioxenophilius* indeed belonged to the xylene degradation pathway (Figure 5) and were detected in equal abundance in both the Coastal and Offshore experiments. However, the total abundance of this pathway was substantially lower than the benzoate degradation pathway, indicating this genus likely catalyzes a secondary “shunt” for PAH degradation intermediates.

### Differential Expression of Alkane and PAH Catabolic Pathways

Several factors are at play when considering the ecological dynamics of alkane and PAH degrading microorganisms in marine environments. First is the relative metabolic complexity of these two processes. The catabolic pathway for alkane degradation is comparatively simple: the activation of an alkane to a fatty alcohol is a single step process through the action of an alkane hydroxylase (44). Once an alkane is activated, the subsequent fatty alcohol quickly feeds through a couple intermediates into the beta-oxidation pathway to produce several acetyl-CoA molecules, a central TCA cycle intermediate. In contrast, PAH degradation involves a complex series of multi-enzyme ring hydroxylation and ring cleavage steps (31). Considering the energetic costs with synthesizing these enzymes, there is likely a fitness cost associated with biodegrading PAHs, especially those of high-molecular weight, compared to biodegrading alkanes. This is consistent with patterns of crude oil weathering typically observed in the environment, where the alkane fraction of crude oil typically disappears faster than the PAH fraction (45–48).

One of the most striking results from our metatranscriptomic datasets was the difference in the expression of alkane and PAH catabolic pathways between the Coastal and Offshore experiments. In particular, the substantially lower expression of alkane activation genes concurrently with increased expression of PAH degradation genes in the Offshore experiment (Figure 5) was unexpected as it contrasts with the above notion that n-alkanes are more labile and are attacked before the more refractory PAHs. This is not due to a lack of alkane degrading taxa—known alkane degrading species such as *Marinobacter* (49), *Thalassolituus* (50), and *Oleibacter* (51) were present in both experiments (Figure 3).

Laboratory studies have estimated that the input of phytoplankton-derived alkane production into marine surface waters is ∼100-fold greater than the combined inputs from oil spills and natural oil seeps (52) and can sustain populations of alkane-degrading bacteria which rapidly expand upon exposure to crude oil (53, 54). Thus, the differences in alkane activation transcripts we observed between the two experiments may be due to a priming effect from the larger abundances of phytoplankton typically observed in Gulf of Mexico coastal shelf waters. Three lines of evidence in our data support this hypothesis: [1] ASVs classified as Cyanobacteria and/or Chloroplasts were substantially more abundant in our Coastal mesocosms than our Offshore mesocosms (Figure 2), [2] transcripts for aldehyde-deformylating oxygenase—the key enzyme in cyanobacterial alkane biosynthesis (55, 56)—were exclusively observed in our coastal metatranscriptomes, and [3] the concentration of naturally present alkanes (i.e. in the Controls) was approximately 10-fold higher in our coastal seawater samples than those collected offshore (Figure 1, Table S2).

With regards to the expression of PAH degradation pathways in our metatranscriptomes, a major difference between the Coastal and Offshore experiment appeared to be due to a differential response of *Cycloclasticus* ecotypes. Our data initially appears to suggest that members of this genus are sensitive to highly concentrated (∼50ppm) CEWAF preparations and instead prefer the lower oil concentrations present in WAF and DCEWAF. We have seen this before in a previous mesocosm experiment we conducted with near-shore seawater: *Cyloclasticus*-related OTUs were nearly completely absent in CEWAF treatments, but thrived in WAF and DCEWAF (37). However, in the current study, this sensitivity enigmatically was not consistent between the Coastal and Offshore experiments. In our metatranscriptomes, the Offshore CEWAF treatment contained the largest abundance of *Cycloclasticus* affiliated dioxygenase and PAH degradation gene transcripts while the Coastal CEWAF treatment contained virtually none. In our 16S rRNA datasets, *Cycloclasticus* related ASVs bloomed from 0.05% to 10.3% relative abundance in the Offshore CEWAF but were absent in the Coastal CEWAF (Figure 3). Although there is evidence that some *Cycloclasticus* species are sensitive to high concentrations of inorganic nutrients (57) or differences in temperature (58), this does not provide a valid explanation for our findings as we amended all mesocosms with f/20 media and both experiments were performed at the same temperature. Taken together, these new findings indicate that a strong selective pressure against *Cycloclasticus* sp. occurred in coastal seawater at high concentrations of chemically dispersed oil, but this negative selection was not solely due to nutrient concentrations, the presence of chemical dispersants, or high concentrations of oil. Further research will be needed to elucidate the cause of this apparent niche partitioning by *Cycloclasticus* ecotypes. Possibilities include variable physiological responses to oil exposure under different environmental regimes, or ecological competition with other microorganisms.

## CONCLUSIONS

In this study, we showed that prokaryotic communities can respond in different ways to oil spills depending on the location of the spill, which is tightly tied to the starting initial community. In our experimental design, season, oil, dispersant, and inorganic nutrients were controlled for between the two experiments. Only community composition and potentially chemical characteristics not measured here were different, indicating that these were responsible for the different responses observed. This is important because oil spills can and do spread out over multiple marine realms (e.g. 2010 Deepwater Horizon, 1979 Ixtoc-1, Gulf War, and 1978 Amoco Cadiz oil spills). The response of microbial communities within those realms may vary and this should weigh into management and mitigation decisions during and after an oil spill. As was shown here, different ecotypes may respond differently to oil and/or dispersant. Gene expression can vary on the community level between different environments facing the same spill. Previous work has shown that niche partitioning may be important to compositional responses to oil spills (7, 37), but this work is the among the first to show that gene expression is also impacted. We conclude that niche partitioning and ecotype dynamics play a critically important role in how marine environments response to past and future oil spills.

## MATERIALS AND METHODS

### Mesocosm Experiments and Sampling

Surface seawater (1m) for the Coastal and Offshore mesocosm experiments were collected at 29°38’ N, 93°50’ W, on July 16th, 2016 and at 29°53’ N, 94°20’ W, on July 9th, 2016, respectively (Figure S1, Table S1). The seawater was transferred to a holding tank in the Texas A&M University at Galveston Sea Life Facility, covered, and stored at room temperature overnight prior to experiment initiation. Four treatments were prepared in triplicate as described previously in Doyle et al. (37): [1] “Control”, containing only seawater, [2] “WAF”, containing seawater and oil supplied as a water-accommodated fraction, [3] “CEWAF”, containing seawater, oil, and Corexit supplied as a chemically-enhanced water-accommodated fraction in a dispersant to oil ratio of 1:20, and [4] “DCEWAF”, a 1:10 dilution of the “CEWAF” treatment. The WAF and CEWAF was prepared by adding 25 mL (5 mL every 30 min for 2.5 h) of unweathered Macondo surrogate oil (WAF) or oil plus dispersant (CEWAF) into 130L of seawater in duplicate baffled recirculation tanks (BRT) and allowing the oil and seawater to mix for ∼24 h (59). Mesocosm tanks were then filled by withdrawing WAF from the bottom of the recirculation tanks in order to avoid including any non-accommodated oil floating on the surface of the BRTs. Each mesocosm tank contained 90L of seawater supplemented at the start of the experiment with 9mL of f/20 nutrient media (N, P, Si) prepared using the Guillard and Ryther (60). This increased inorganic nitrogen concentrations by approximately 2.7-fold and 3.8-fold, respectively, in the Coastal and Offshore experiments. Likewise, silicate and inorganic phosphate concentrations were increased by approximately 1.5-fold in both experiments. This was done to ensure microbial communities would not be nutrient limited during the experiment. In order to minimize potential bottle effects, 90L mesocosms were chosen so that no more than 10% of the total volume was removed by sampling during the course of the experiment. Full-spectrum fluorescent lamps (UV-Vis 375-750nm; Sylvania GRO-LUX; Wilmington, MA, USA) provided a 12h light/12h dark cycle (50-80 µmol photons m^-2^ s^-1^) and the room was kept at ∼21°C.

T0 sampling for each experiment began immediately after the generation of the WAF, DCEWAF, and CEWAF treatments was complete, corresponding to 24 hours after oil addition. Each experiment was run until the remaining oil concentration within the CEWAF treatment (highest oil concentration) reached ∼20% of the initial oil concentration (61, 62). This was 72h and 96h for the Coastal and Offshore experiments, respectively. For each experiment, starting at time zero and every 12h thereafter, ∼1L of water was collected from each mesocosm in a clean, opaque Nalgene bottle through a PTFE-lined spigot mounted on the side of each tank (10cm above bottom). For cell count samples, 10mL of this water was fixed with formalin (final concentration 2%) and stored at 4°C. For DNA samples, the collected water was prefiltered through a 10µm filter to exclude zooplankton and large eukaryotic cells followed by filtering 200mL onto a 47mm, 0.22µm Supor PES membrane (Pall; Port Washington, NY, USA). For RNA samples, up to 800mL (or until clogged) of the remaining 10µm-prefiltered water was filtered onto 47mm, 0.22µm Supor PES membranes. After collection, all filters were placed in cryotubes and stored at -80°C. RNA filters were immersed in 1mL of TE buffer (10mM Tris, 1mM EDTA, pH 7.5) before freezing.

### Analysis of hydrocarbon chemistry

Samples (1-3.5 L) were collected every 24 hours in amber bottles with Teflon lined screw caps from each of the triplicate treatment tanks and immediately amended with ∼20mL of dichloromethane (DCM). Prior to extraction, PAH surrogates (d8-naphthalene, d10-acenaphthene, d10-phenanthrene, d12-chrysene, and d12-perylene) and aliphatic surrogate standards (deuterated nC12, nC20, nC24, and nC30) were added (63). The DCM mixture was reduced, exchanged into hexane, and then transferred to silica gel/alumina columns for purification (59). Hydrocarbons were eluted with 200 ml of a 1:1 pentane/DCM solution, evaporated, and exchanged with hexane (64, 65). Aliphatic hydrocarbons were then analyzed on an Agilent 7890 gas chromatograph with a flame ionization detector (GC/FID) according to Wade et al. (64) with updates in Morales-McDevitt et al. (63). PAHs were analyzed on a Hewlett-Packard 6890 gas chromatograph coupled with a Hewlett-Packard 5973 mass selective detector. A laboratory reference sample was analyzed with each batch of samples to confirm GC/MS/SIM system performance and calibration (64, 65). Alkylated PAH were quantitated based response of the parent PAH compound (63).

### Cell Abundance

Formalin preserved samples were stained with DAPI (45µM final concentration), filtered onto black polycarbonate filters (25mm, 0.2µm), mounted on a glass microscope slide with two drops of Citifluor AF1 antifade, and directly counted with an epifluorescence microscope (Axio Imager M2, Zeiss; Jena, Germany). A minimum of 10 fields were randomly counted per sample.

### 16S rRNA Gene Amplicon Sequencing

The V4 hyper-variable region of the 16S rRNA gene was amplified from each sample (n=190) following the protocol previously described in Doyle et al. (37). Briefly, filters were sliced into small pieces using a sterile scalpel and subsequently extracted using FastDNA Spin kits (MP Biomedical; Santa Ana, CA, USA). Five sample-free filters were processed as protocol blanks. Universal (Bacteria and Archaea) V4 primers 515F and 806R primers were used for amplification (66) . Sequencing was performed on the Illumina MiSeq platform (500 cycle, V2 chemistry) at the Georgia Genomics Facility (Athens, GA, USA). Raw read curation and processing into ASVs was performed by following the standard pipeline of the DADA2 package (67) in R (details below; Figure S10). All ASV tables were subsampled without replacement to an even depth (n=7754, minimum value of samples, mean ∼60,000; Table S6) before downstream ecological analyses were performed with a combination of mothur v1.42.1, phyloseq v1.28, and/or vegan v.2.5-6 (68–70).

### 16S rRNA Amplicon Sequence Analysis

Raw reads were processed in DADA2 v.1.12.1 using standard filtering parameters (maxN=0, truncQ=2, rm.phix=TRUE, and maxEE=2). Quality profiles of the forward (R1) and reverse (R2) reads were manually inspected and then reads were truncated to the length after which the distribution of quality scores began to drop: 240bp and 160bp, respectively. Error rates for the filtered and trimmed R1 and R2 reads were calculated using the learnErrors function and subsequently used to denoise reads using the DADA2 sample inference algorithm. The denoised R1 and R2 reads, free of substitution and indel errors, were then merged together into amplicon sequence variants (ASV) using a global ends-free alignment. Paired reads containing any mismatches in the overlapping region were removed from the dataset. Chimeric ASVs were identified and removed by using the consensus method within the removeBimeraDenovo function. The number of reads that made it through each step in the pipeline for each sample is detailed in Table S7. As a final curation step, any ASVs of which ≥0.1% of its reads were from one of the protocol blanks were removed. A total of 11,654,656 sequences passed our quality control steps, corresponding to an average of 59,716 sequences per sample, and were used to construct a curated library containing 2880 ASVs (Table S4). Rarefaction curves for all samples indicated that any unsampled diversity contained only rare members (Figure S10). A consensus taxonomy for each ASV was assigned using the naïve Bayesian classified method (71) trained on release 128 of the SILVA reference database (72).

### Redundancy Analysis

Redundancy analyses (RDA) were performed to investigate the relationship between microbial community compositions, hours of incubation, and initial oil concentrations and to identify ASVs associated with the presence of oil. For these analyses, ASVs without an average relative abundance ≥0.2%, after subsampling, in at least one sample were excluded. ASV counts were transformed using using a Hellinger transformation and an RDA model was calculated using hours and initial oil concentration as variables (73). The significance of each RDA model was assessed using a permutational ANOVA test (74) and the variance explained by each variable was estimated by variance partitioning with partial RDAs (75). ASVs associated with the presence of oil were identified as those whose RDA-weighted average loading scores were positively correlated with the presence of oil and were greater than the equilibrium contribution of the RDA model (the proportion of variance which would be explained by a random canonical axis).

### RNA Extraction and Metatranscriptome Sequencing

For each mesocosm experiment, three time points, corresponding to the start, middle, and end of the experiment, were selected for metatranscriptomic sequencing (0h, 36h, and 72h for the Coastal experiment, and 0h, 48h, and 96h for the Offshore experiment). To ensure an adequate quantity of RNA was recovered for sequencing, the triplicate samples for each treatment were combined into a single extraction. A customized phenol:chloroform RNA extraction was used to isolate RNA. Before thawing samples for RNA extraction, β-mercaptoethanol was added to each sample to a final concentration of 1% (v/v). Once thawed, filters were sliced into small pieces using a sterile scalpel and placed into a bead-beating tube along with the TE buffer (1mM EDTA, 10mM Tris; pH 6.3) in which they were frozen and ∼1g of sterilized 0.1mm diameter zirconia/silica beads (BioSpec Products; Bartlesville, OK, USA). Samples were then homogenized in a BioSpec Mini-Beadbeater for 2 minutes at maximum speed. After bead beating, crude extracts were amended with two volumes of chilled (4°C) denaturing buffer (4M guanidine thiocyanate, 50mM Tris, 10mM EDTA, 1% w/v N-lauroylsarcosine, 1% β-mercaptoethanol). Insoluble material was pelleted via centrifugation (4500×g for 5 min at 4°C) and the supernatant was collected. The pellet was washed with 3mL of chilled denaturing buffer, centrifuged again, and the resulting supernatant pooled with the first. This pooled lysate was then extracted with an equal volume of phenol:chloroform:isoamyl alcohol (25:24:1, pH 6.6), followed by a second extraction with chloroform:isoamyl alcohol (24:1). Nucleic acids were purified from these extracts via an overnight isopropanol precipitation with 3M sodium acetate (pH 6.0) and a subsequent 70% ethanol wash, followed by resuspension in 100µL of TE buffer. Genomic DNA was eliminated from RNA samples with TURBO DNA-free kits (Ambion; Waltham, MA, USA) followed by purification with MEGAclear Transcription Clean-up kits (Ambion). The resulting purified total RNA extracts were processed with MICROBExpress kits (Ambion) to reduce the amount of 16S and 23S rRNA transcripts within the samples and then sent to the University of Delaware DNA Sequencing & Genotyping Center (Newark, DE, USA) for Illumina HiSeq 2500 sequencing (paired-end 150bp reads).

We obtained 576 million paired reads across 24 samples from RNA sequencing. Ribosomal RNA (rRNA) reads were filtered from the dataset using SortMeRNA v2.1 (76) with all eight prepackaged rRNA reference databases, and an E-value threshold of 1e-20 (Table S8). The resulting rRNA-depleted paired end reads were quality filtered and trimmed with Trimmomatic v0.36 (77) to remove Illumina adapters and low-quality base pairs with the following parameters: Sliding Window:4:5, Headcrop:10, Leading:5, Trailing:5, Minimum Length:115. Successful removal of adapter sequences and low-quality reads was confirmed with FastQC v0.11.9 (78).

### Metatranscriptome analysis

De novo transcriptome assembly was performed with the curated paired reads using Trinity v2.5.1 (79) with default parameters, producing 648 351 contigs ranging between 201 and 54 422 bp in length. Contig abundances within each sample were then quantified by mapping reads to the metatranscriptome assembly with kallisto v0.44.0 (80). Read counts were normalized for contig length into TPM (transcripts per million transcripts) values as previously described (81), and scaled for cross-sample comparison using the trimmed mean of M-values (TMM) method described by Robinson and Oshlack (82). Open reading frames were identified and translated into amino acid sequences using Prodigal v2.6.3 in anonymous mode (83). Functional annotations were performed using a GHOSTX v1.3.7 search against the Kyoto Encyclopedia of Genes and Genomes (KEGG) database using the single-directional best hit assignment method targeted to prokaryotes. A custom R script was then used to select all transcripts whose functional annotation belonged to a KEGG pathway involved in the degradation of saturated hydrocarbons (alkanes and cycloalkanes), PAHs, or their metabolic intermediates to central metabolism (glycolysis, TCA cycle) substrates (Table S9). We also searched for genes involved in the initial oxidation of alkanes using a BLASTP (single best hit ≤1E^-20^, alignment length ≥100, bit score ≥50) search of a custom database (Table S10) of soluble cytochrome P450 alkane hydroxylases of the CYP153 family (49). These proteins would have been missed from the above KEGG-based analysis as there are currently no KEGG orthologs specific to this family of enzymes (84). Identified hydrocarbon degradation gene transcripts were taxonomically classified using KAIJU v1.6.3 in greedy mode (5 substitutions allowed) with the NCBI non-redundant database (+eukaryotes) as a reference (85). Transcripts with identically scored matches to different taxa were classified to the least common ancestor in the phylogenetic tree. Following the method described in Jenior et al. (86), an outlier analysis was performed to estimate the number of highly expressed transcripts within each sample.

## ACKNOWLEDGMENTS

Data is publicly available through the Gulf of Mexico Research Initiative Information and Data Cooperative (GRIIDC) at http://data.gulfresearchinitative.org DOIs: 10.7266/N77D2SPZ (Coastal 16S rRNA libraries), 10.7266/N7C53JDP (Offshore 16S rRNA libraries), 10.7266/9EDJRA3Q (metatranscriptome sequences), 10.7266/N74X568X (Coastal alkane and PAH measurements), and 10.7266/N78P5XZD (Offshore alkane and PAH measurements).

---

Figure S1. Map showing the locations where seawater for the Coastal and Offshore experiments were collected.

Figure S2. Observed cell abundances with each treatment of the Coastal and Offshore experiments. Error bars denote standard deviations.

Figure S3. NMDS ordination of Bray-Curtis dissimilarities between samples from both experiments. In this case, water location (i.e. coastal vs offshore) dominates the sample clustering within the ordination. Numbered labels indicate hours of incubation.

Figure S4. (A, E) NMDS ordination of Bray-Curtis dissimilarities in the Offshore and Coastal experiments indicated microbial community structure was driven by the level of oil exposure (treatment). Numbered labels indicate hours of incubation. (B-D and F-H) RDA score triplots for the three oil-amended treatments within the Offshore and Coastal experiments, respectively. Each treatment was modeled against the Control. Colored circles represent samples. Plus (+) signs represent ASVs. Purple rings represent the equilibrium contribution of the overall model. Arrows represent the vectors of the two explanatory variables used in the model: (1) the presence of oil, and (2) hours of incubation. Right-angled scalar projections of an ASV point onto the oil vector approximates that ASV’s association with the presence of oil. Bolded, red ASVs were designated as oil-associated for this study.

Figure S5. Relative abundances of abundant microbial lineages observed in each experiment based on the metatranscriptomes. The plot was built to display the highest resolution classification for the most abundant taxa. The plot was constructed as follows: First, transcripts were clustered at the genus level and any genera having a relative abundance ≥20% in at least one of the samples were plotted. This procedure was subsequently repeated with the remaining unplotted transcripts at the taxonomic levels of family, order, class and phylum. Any remaining rare transcripts left after this procedure were not plotted.

Figure S6. Observed variation in the relative abundance of the 10 most abundant orders in the 16S rRNA amplicon dataset. Boxplots display variation between time points for each treatment. Cross-lines indicated the median observed relative abundance. Larger boxes indicate increased variability over time within that respective treatment.

Figure S7. Outlier analysis of hydrocarbon degradation gene expression within the Coastal WAF, DCEWAF, and CEWAF treatments versus the Control treatments. Each point represents a unique open reading frame identified in the respective metatranscriptomes. Outliers were defined using a squared residual cutoff value of 2 versus the x = y dotted-line. Points are colored by taxonomic assignment (genus) of each transcript. Gray points denote those genes whose abundance were comparable between the respective oil-amended treatment and the Control. Outliers above the shaded-gray ribbon were considered highly expressed. Outlier analysis was performed using a modification of the custom R-scripts described in Jenior et al. (2018).

Figure S8. Outlier analysis of hydrocarbon degradation gene expression within the Offshore WAF, DCEWAF, and CEWAF treatments versus the Control treatments. Each point represents a unique open reading frame identified in the respective metatranscriptomes. Outliers were defined using a squared residual cutoff value of 2 versus the x = y dotted-line. Points are colored by taxonomic assignment (genus) of each transcript. Gray points denote those genes whose abundance were comparable between the respective oil-amended treatment and the Control. Outliers above the shaded-gray ribbon were considered highly expressed. Outlier analysis was performed using a modification of the custom R-scripts described in Jenior et al. (2018).

Figure S9. Hydrocarbon degradation genes were identified by comparing their abundance in each oil-amended treatment with that observed in the respective Control. Although the number of upregulated genes differed between treatments (see Figure S6 and S6), the amount those genes were upregulated did not differ between treatment.

Figure S10. Rarefaction curves for all 16S ASV libraries reached saturation.

Table S1. Seawater samples collected for mesocosm experiments.

Table S2. Measured concentrations (µg/L) of total alkanes and total PAHs over time in both mesocosm experiments. Half-lifes (t0.5) were calculated over 72 hours. %RSD denotes the relative standard deviation. n.d. indicates half-life not determined.

Table S3. Linear regression models of cell abundances over time.

Table S4. Read count within each sample, taxonomy, and sequence of each ASV.

Table S5. Pairwise nucleotide position differences and percent identity in the 16S rRNA gene (V4 hypervariable region) of oil-enriched ASVs belonging to the same genus.

Table S6. Alpha Diversity Summary of 16S rRNA amplicon libraries. (* denotes values calculated after normalizing read-depth by subsampling each library to 7754 reads; 1000 iterations)

Table S7. Number of sequence reads that made it through each step of the DADA2 pipeline.

Table S8. Metatranscriptome reads identified as rRNA using SortMeRNA.

Table S9. All transcripts whose KEGG annotation, generated using GHOSTX, matched one of the following were identified as hydrocarbon degradation gene transcripts.

Table S10. List of proteins in our CYP153 database. We used this database along with BLASTP to identify transcripts for P450 alkane hydroxylases of the CYP153 family.

## REFERENCES

1. Tissot BP, Welte DH. 1984. Petroleum formation and occurrence, 2nd ed. Springer: Berlin, Heidelberg, Germany.

2. Head IM, Jones DM, Röling WFM. 2006. Marine microorganisms make a meal of oil. Nature Reviews Microbiology 4:173–182.

3. Lessard RR, DeMarco G. 2000. The Significance of Oil Spill Dispersants. Spill Science & Technology Bulletin 6:59–68.

4. Brakstad OG, Ribicic D, Winkler A, Netzer R. 2018. Biodegradation of dispersed oil in seawater is not inhibited by a commercial oil spill dispersant. Marine Pollution Bulletin 129:555–561.

5. Hamdan L, Fulmer P. 2011. Effects of COREXIT® EC9500A on bacteria from a beach oiled by the Deepwater Horizon spill. Aquatic Microbial Ecology 63:101–109.

6. Kleindienst S, Seidel M, Ziervogel K, Grim S, Loftis K, Harrison S, Malkin SY, Perkins MJ, Field J, Sogin ML, Dittmar T, Passow U, Medeiros PM, Joye SB. 2015. Chemical dispersants can suppress the activity of natural oil-degrading microorganisms. Proceedings of the National Academy of Sciences 112:14900–14905.

7. Techtmann SM, Zhuang M, Campo P, Holder E, Elk M, Hazen TC, Conmy R, Domingo JWS. 2017. Corexit 9500 Enhances Oil Biodegradation and Changes Active Bacterial Community Structure of Oil-Enriched Microcosms. Appl Environ Microbiol 83:e03462–16.

8. Tremblay J, Yergeau E, Fortin N, Cobanli S, Elias M, King TL, Lee K, Greer CW. 2017. Chemical dispersants enhance the activity of oil-and gas condensate-degrading marine bacteria. The ISME Journal.

9. Atlas RM, Hazen TC. 2011. Oil Biodegradation and Bioremediation: A Tale of the Two Worst Spills in U.S. History. Environmental Science & Technology 45:6709–6715.

10. Kujawinski EB, Kido Soule MC, Valentine DL, Boysen AK, Longnecker K, Redmond MC. 2011. Fate of Dispersants Associated with the Deepwater Horizon Oil Spill. Environ Sci Technol 45:1298–1306.

11. Dubinsky EA, Conrad ME, Chakraborty R, Bill M, Borglin SE, Hollibaugh JT, Mason OU, M. Piceno Y, Reid FC, Stringfellow WT, Tom LM, Hazen TC, Andersen GL. 2013. Succession of Hydrocarbon-Degrading Bacteria in the Aftermath of the *Deepwater Horizon* Oil Spill in the Gulf of Mexico. Environmental Science & Technology 47:10860–10867.

12. Gutierrez T, Singleton DR, Berry D, Yang T, Aitken MD, Teske A. 2013. Hydrocarbon-degrading bacteria enriched by the Deepwater Horizon oil spill identified by cultivation and DNA-SIP. The ISME journal 7:2091.

13. Hazen TC, Dubinsky EA, DeSantis TZ, Andersen GL, Piceno YM, Singh N, Jansson JK, Probst A, Borglin SE, Fortney JL, Stringfellow WT, Bill M, Conrad ME, Tom LM, Chavarria KL, Alusi TR, Lamendella R, Joyner DC, Spier C, Baelum J, Auer M, Zemla ML, Chakraborty R, Sonnenthal EL, D’haeseleer P, Holman H-YN, Osman S, Lu Z, Nostrand JDV, Deng Y, Zhou J, Mason OU. 2010. Deep-Sea Oil Plume Enriches Indigenous Oil-Degrading Bacteria. Science 330:204–208.

14. Mason OU, Hazen TC, Borglin S, Chain PS, Dubinsky EA, Fortney JL, Han J, Holman H- YN, Hultman J, Lamendella R, others. 2012. Metagenome, metatranscriptome and single-cell sequencing reveal microbial response to Deepwater Horizon oil spill. The ISME journal 6:1715–1727.

15. Rivers AR, Sharma S, Tringe SG, Martin J, Joye SB, Moran MA. 2013. Transcriptional response of bathypelagic marine bacterioplankton to the Deepwater Horizon oil spill. The ISME journal 7:2315–2329.

16. Engel AS, Gupta AA. 2014. Regime Shift in Sandy Beach Microbial Communities following Deepwater Horizon Oil Spill Remediation Efforts. PLoS ONE 9:e102934.

17. Kostka JE, Prakash O, Overholt WA, Green SJ, Freyer G, Canion A, Delgardio J, Norton N, Hazen TC, Huettel M. 2011. Hydrocarbon-Degrading Bacteria and the Bacterial Community Response in Gulf of Mexico Beach Sands Impacted by the Deepwater Horizon Oil Spill. Applied and Environmental Microbiology 77:7962–7974.

18. Lamendella R, Strutt S, Borglin SE, Chakraborty R, Tas N, Mason OU, Hultman J, Prestat E, Hazen TC, Jansson J. 2014. Assessment of the Deepwater Horizon oil spill impact on Gulf coast microbial communities. Front Microbiol 5.

19. Rodriguez-r LM, Overholt WA, Hagan C, Huettel M, Kostka JE, Konstantinidis KT. 2015. Microbial community successional patterns in beach sands impacted by the Deepwater Horizon oil spill. The ISME journal.

20. Beazley MJ, Martinez RJ, Rajan S, Powell J, Piceno YM, Tom LM, Andersen GL, Hazen TC, Van Nostrand JD, Zhou J, Mortazavi B, Sobecky PA. 2012. Microbial Community Analysis of a Coastal Salt Marsh Affected by the Deepwater Horizon Oil Spill. PLoS ONE 7:e41305.

21. Mahmoudi N, Porter TM, Zimmerman AR, Fulthorpe RR, Kasozi GN, Silliman BR, Slater GF. 2013. Rapid Degradation of Deepwater Horizon Spilled Oil by Indigenous Microbial Communities in Louisiana Saltmarsh Sediments. Environ Sci Technol 47:13303–13312.

22. King GM, Kostka JE, Hazen TC, Sobecky PA. 2015. Microbial Responses to the *Deepwater Horizon* Oil Spill: From Coastal Wetlands to the Deep Sea. Annual Review of Marine Science 7:377–401.

23. Fortunato CS, Crump BC. 2011. Bacterioplankton Community Variation Across River to Ocean Environmental Gradients. Microb Ecol 62:374–382.

24. D’Ambrosio L, Ziervogel K, MacGregor B, Teske A, Arnosti C. 2014. Composition and enzymatic function of particle-associated and free-living bacteria: a coastal/offshore comparison. The ISME Journal 8:2167–2179.

25. Lee K, Nedwed T, Prince RC, Palandro D. 2013. Lab tests on the biodegradation of chemically dispersed oil should consider the rapid dilution that occurs at sea. Marine Pollution Bulletin 73:314–318.

26. Watson SW, Novitsky TJ, Quinby HL, Valois FW. 1977. Determination of bacterial number and biomass in the marine environment. Applied and Environmental Microbiology 33:940–946.

27. Bianchi A, Giuliano L. 1996. Enumeration of viable bacteria in the marine pelagic environment. Applied and environmental microbiology 62:174–177.

28. Gutierrez T, Green DH, Nichols PD, Whitman WB, Semple KT, Aitken MD. 2013. Polycyclovorans algicola gen. nov., sp. nov., an Aromatic-Hydrocarbon-Degrading Marine Bacterium Found Associated with Laboratory Cultures of Marine Phytoplankton. Applied and Environmental Microbiology 79:205–214.

29. Van Beilen JB, Li Z, Duetz WA, Smits THM, Witholt B. 2003. Diversity of Alkane Hydroxylase Systems in the Environment. Oil & Gas Science and Technology 58:427– 440.

30. Johnsen AR, Wick LY, Harms H. 2005. Principles of microbial PAH-degradation in soil. Environmental Pollution 133:71–84.

31. Seo J-S, Keum Y-S, Li QX. 2009. Bacterial Degradation of Aromatic Compounds. Int J Environ Res Public Health 6:278–309.

32. Fulco AJ. 1983. Fatty acid metabolism in bacteria. Progress in Lipid Research 22:133– 160.

33. Dyksterhouse SE, Gray JP, Herwig RP, Lara JC, Staley JT. 1995. Cycloclasticus pugetii gen. nov., sp. nov., an aromatic hydrocarbon-degrading bacterium from marine sediments. International Journal of Systematic and Evolutionary Microbiology 45:116– 123.

34. Yakimov MM, Timmis KN, Golyshin PN. 2007. Obligate oil-degrading marine bacteria. Current Opinion in Biotechnology 18:257–266.

35. Orcutt BN, Joye SB, Kleindienst S, Knittel K, Ramette A, Reitz A, Samarkin V, Treude T, Boetius A. 2010. Impact of natural oil and higher hydrocarbons on microbial diversity, distribution, and activity in Gulf of Mexico cold-seep sediments. Deep Sea Research Part II: Topical Studies in Oceanography 57:2008–2021.

36. Sun X, Kostka JE. 2019. Hydrocarbon-Degrading Microbial Communities Are Site Specific, and Their Activity Is Limited by Synergies in Temperature and Nutrient Availability in Surface Ocean Waters. Appl Environ Microbiol 85.

37. Doyle SM, Whitaker EA, De Pascuale V, Wade TL, Knap AH, Santschi PH, Quigg A, Sylvan JB. 2018. Rapid Formation of Microbe-Oil Aggregates and Changes in Community Composition in Coastal Surface Water Following Exposure to Oil and the Dispersant Corexit. Front Microbiol 9.

38. Redmond MC, Valentine DL. 2012. Natural gas and temperature structured a microbial community response to the Deepwater Horizon oil spill. Proceedings of the National Academy of Sciences 109:20292–20297.

39. Wang W, Zhong R, Shan D, Shao Z. 2014. Indigenous oil-degrading bacteria in crude oil-contaminated seawater of the Yellow sea, China. Applied Microbiology and Biotechnology 98:7253–7269.

40. Jin HM, Kim JM, Lee HJ, Madsen EL, Jeon CO. 2012. Alteromonas As a Key Agent of Polycyclic Aromatic Hydrocarbon Biodegradation in Crude Oil-Contaminated Coastal Sediment. Environ Sci Technol 46:7731–7740.

41. McGenity TJ, Folwell BD, McKew BA, Sanni GO. 2012. Marine crude-oil biodegradation: a central role for interspecies interactions. Aquatic Biosystems 8:10.

42. Stoddard SF, Smith BJ, Hein R, Roller BRK, Schmidt TM. 2015. rrnDB: improved tools for interpreting rRNA gene abundance in bacteria and archaea and a new foundation for future development. Nucleic Acids Research 43:D593–D598.

43. Iwaki H, Yamamoto T, Hasegawa Y. 2018. Isolation of marine xylene-utilizing bacteria and characterization of Halioxenophilus aromaticivorans gen. nov., sp. nov. and its xylene degradation gene cluster. FEMS Microbiol Lett 365.

44. Rojo F. 2009. Degradation of alkanes by bacteria. Environmental Microbiology 11:2477–2490.

45. Das N, Chandran P. 2011. Microbial Degradation of Petroleum Hydrocarbon Contaminants: An Overview. Biotechnology Research International 2011:1–13.

46. Sugiura K, Ishihara M, Shimauchi T, Harayama S. 1997. Physicochemical Properties and Biodegradability of Crude Oil. Environ Sci Technol 31:45–51.

47. Wang Z, Fingas M, Blenkinsopp S, Sergy G, Landriault M, Sigouin L, Foght J, Semple K, Westlake DWS. 1998. Comparison of oil composition changes due to biodegradation and physical weathering in different oils. Journal of Chromatography A 809:89–107.

48. Jonker MTO, Brils JM, Sinke AJC, Murk AJ, Koelmans AA. 2006. Weathering and toxicity of marine sediments contaminated with oils and polycyclic aromatic hydrocarbons. Environmental Toxicology and Chemistry 25:1345–1353.

49. Nie Y, Chi C-Q, Fang H, Liang J-L, Lu S-L, Lai G-L, Tang Y-Q, Wu X-L. 2014. Diverse alkane hydroxylase genes in microorganisms and environments. Scientific Reports 4:4968.

50. Yakimov MM. 2004. Thalassolituus oleivorans gen. nov., sp. nov., a novel marine bacterium that obligately utilizes hydrocarbons. INTERNATIONAL JOURNAL OF SYSTEMATIC AND EVOLUTIONARY MICROBIOLOGY 54:141–148.

51. Teramoto M, Ohuchi M, Hatmanti A, Darmayati Y, Widyastuti Y, Harayama S, Fukunaga Y. 2011. Oleibacter marinus gen. nov., sp. nov., a bacterium that degrades petroleum aliphatic hydrocarbons in a tropical marine environment. International Journal of Systematic and Evolutionary Microbiology 61:375–380.

52. 52. White HK, Marx CT, Valentine DL, Sharpless CM, Aeppli C, Gosselin K, Kivenson V, Liu RM, Nelson RK, Sylva SP, Reddy CM. 2019. Examining inputs of biogenic and oil-derived hydrocarbons in surface waters following the Deepwater Horizon oil spill. ACS Earth Space Chem acsearthspacechem.9b00090.

53. Lea-Smith DJ, Biller SJ, Davey MP, Cotton CAR, Sepulveda BMP, Turchyn AV, Scanlan DJ, Smith AG, Chisholm SW, Howe CJ. 2015. Contribution of cyanobacterial alkane production to the ocean hydrocarbon cycle. PNAS 112:13591–13596.

54. Valentine DL, Reddy CM. 2015. Latent hydrocarbons from cyanobacteria. PNAS 112:13434–13435.

55. Li N, Chang W, Warui DM, Booker SJ, Krebs C, Bollinger JM. 2012. Evidence for Only Oxygenative Cleavage of Aldehydes to Alk(a/e)nes and Formate by Cyanobacterial Aldehyde Decarbonylases. Biochemistry 51:7908–7916.

56. Jiménez-Díaz, L, Caballero A, Pérez-Hernández N, Segura A. 2017. Microbial alkane production for jet fuel industry: motivation, state of the art and perspectives. Microbial Biotechnology 10:103–124.

57. Singh AK, Sherry A, Gray ND, Jones DM, Bowler BFJ, Head IM. 2014. Kinetic parameters for nutrient enhanced crude oil biodegradation in intertidal marine sediments. Front Microbiol 5.

58. Teira E, Lekunberri I, Gasol JM, Nieto-Cid M, Álvarez-Salgado XA, Figueiras FG. 2007. Dynamics of the hydrocarbon-degrading Cycloclasticus bacteria during mesocosm-simulated oil spills. Environmental Microbiology 9:2551–2562.

59. Knap AH, Sleeter TD, Dodge RE, Wyers SC, Frith HR, Smith SR. 1983. The effects of oil spills and dispersant use on corals: a review and multidisciplinary experimental approach. Oil and Petrochemical Pollution 1:157–169.

60. Guillard RRL, Ryther JH. 1962. STUDIES OF MARINE PLANKTONIC DIATOMS: I. CYCLOTELLA NANA HUSTEDT, AND DETONULA CONFERVACEA (CLEVE) GRAN. Canadian Journal of Microbiology 8:229–239.

61. Kamalanathan M, Xu C, Schwehr K, Bretherton L, Beaver M, Doyle SM, Genzer J, Hillhouse J, Sylvan JB, Santschi P, Quigg A. 2018. Extracellular Enzyme Activity Profile in a Chemically Enhanced Water Accommodated Fraction of Surrogate Oil: Toward Understanding Microbial Activities After the Deepwater Horizon Oil Spill. Front Microbiol 9.

62. Xu C, Lin P, Zhang S, Sun L, Xing W, Schwehr KA, Chin W-C, Wade TL, Knap AH, Hatcher PG, Yard A, Jiang C, Quigg A, Santschi PH. 2019. The interplay of extracellular polymeric substances and oil/Corexit to affect the petroleum incorporation into sinking marine oil snow in four mesocosms. Science of The Total Environment 693:133626.

63. Morales-McDevitt M, Shi D, Knap AH, Quigg A, Sweet ST, Sericano JL, Wade TL. in review. Fate and transport of petroleum hydrocarbons in controlled mesocosm studies. PLOS ONE.

64. Wade TL, Sweet ST, Sericano JL, Guinasso NL, Diercks A-R, Highsmith RC, Asper VL, Joung D, Shiller AM, Lohrenz SE, Joye SB. 2011. Analyses of Water Samples From the Deepwater Horizon Oil Spill: Documentation of the Subsurface Plume, p. 77–82. In Liu, Y, MacFadyen, A, Ji, Z-G, Weisberg, RH (eds.), Geophysical Monograph Series. American Geophysical Union, Washington, D. C.

65. Wade TL, Sericano JL, Sweet ST, Knap AH, Guinasso NL. 2016. Spatial and temporal distribution of water column total polycyclic aromatic hydrocarbons (PAH) and total petroleum hydrocarbons (TPH) from the Deepwater Horizon (Macondo) incident. Marine Pollution Bulletin 103:286–293.

66. Walters W, Hyde ER, Berg-Lyons D, Ackermann G, Humphrey G, Parada A, Gilbert JA, Jansson JK, Caporaso JG, Fuhrman JA, others. 2016. Improved bacterial 16S rRNA gene (V4 and V4-5) and fungal internal transcribed spacer marker gene primers for microbial community surveys. mSystems 1:e00009–15.

67. Callahan BJ, McMurdie PJ, Rosen MJ, Han AW, Johnson AJA, Holmes SP. 2016. DADA2: High-resolution sample inference from Illumina amplicon data. Nature Methods 13:581–583.

68. McMurdie PJ, Holmes S. 2013. phyloseq: An R Package for Reproducible Interactive Analysis and Graphics of Microbiome Census Data. PLOS ONE 8:e61217.

69. Oksanen J, Blanchet FG, Friendly M, Kindt R, Legendre P, McGlinn D, Minchin PR, O’Hara RB, Simpson GL, Solymos P, Stevens MHH, Szoecs E, Wagner H. 2018. vegan: Community Ecology Package.

70. Schloss PD, Westcott SL, Ryabin T, Hall JR, Hartmann M, Hollister EB, Lesniewski RA, Oakley BB, Parks DH, Robinson CJ, Sahl JW, Stres B, Thallinger GG, Horn DJV, Weber CF. 2009. Introducing mothur: Open-Source, Platform-Independent, Community-Supported Software for Describing and Comparing Microbial Communities. Appl Environ Microbiol 75:7537–7541.

71. Wang Q, Garrity GM, Tiedje JM, Cole JR. 2007. Naïve Bayesian Classifier for Rapid Assignment of rRNA Sequences into the New Bacterial Taxonomy. Appl Environ Microbiol 73:5261–5267.

72. Quast C, Pruesse E, Yilmaz P, Gerken J, Schweer T, Yarza P, Peplies J, Glockner FO. 2013. The SILVA ribosomal RNA gene database project: improved data processing and web-based tools. Nucleic Acids Research 41:D590–D596.

73. Legendre P, Gallagher ED. 2001. Ecologically meaningful transformations for ordination of species data. Oecologia 129:271–280.

74. Legendre P, Oksanen J, Braak CJF ter. 2011. Testing the significance of canonical axes in redundancy analysis. Methods in Ecology and Evolution 2:269–277.

75. Legendre P, Borcard D, Roberts DW. 2012. Variation partitioning involving orthogonal spatial eigenfunction submodels. Ecology 93:1234–1240.

76. Kopylova E, Noé L, Touzet H. 2012. SortMeRNA: fast and accurate filtering of ribosomal RNAs in metatranscriptomic data. Bioinformatics 28:3211–3217.

77. Bolger AM, Lohse M, Usadel B. 2014. Trimmomatic: a flexible trimmer for Illumina sequence data. Bioinformatics 30:2114–2120.

78. Andrews S. 2010. FastQC: a quality control tool for high throughput sequence data.

79. Grabherr MG, Haas BJ, Yassour M, Levin JZ, Thompson DA, Amit I, Adiconis X, Fan L, Raychowdhury R, Zeng Q, Chen Z, Mauceli E, Hacohen N, Gnirke A, Rhind N, Palma F di, Birren BW, Nusbaum C, Lindblad-Toh K, Friedman N, Regev A. 2011. Full-length transcriptome assembly from RNA-Seq data without a reference genome. Nature Biotechnology 29:644–652.

80. Bray NL, Pimentel H, Melsted P, Pachter L. 2016. Near-optimal probabilistic RNA-seq quantification. Nature Biotechnology 34:525–527.

81. Wagner GP, Kin K, Lynch VJ. 2012. Measurement of mRNA abundance using RNA-seq data: RPKM measure is inconsistent among samples. Theory Biosci 131:281–285.

82. Robinson MD, Oshlack A. 2010. A scaling normalization method for differential expression analysis of RNA-seq data. Genome Biology 11:R25.

83. Hyatt D, Chen G-L, LoCascio PF, Land ML, Larimer FW, Hauser LJ. 2010. Prodigal: prokaryotic gene recognition and translation initiation site identification. BMC Bioinformatics 11:119.

84. Tremblay J, Greer C. 2019. Metagenomic Data Mining in Oil Spill Studies, p. 211–223. In Comprehensive Biotechnology 3rd Edition. Pergamon.

85. Menzel P, Ng KL, Krogh A. 2016. Fast and sensitive taxonomic classification for metagenomics with Kaiju. Nature Communications 7:11257.

86. Jenior ML, Leslie JL, Young VB, Schloss PD. 2018. *Clostridium difficile* Alters the Structure and Metabolism of Distinct Cecal Microbiomes during Initial Infection To Promote Sustained Colonization. mSphere 3:e00261–18, /msphere/3/3/mSphere261-18.atom.

